# Crops and the seed mass-seed output trade-off in plants

**DOI:** 10.1101/466250

**Authors:** Adam R. Martin

## Abstract

A trade-off between seed mass (SM) and seed output (SO) defines a central axis of ecological variation among plants, with implications for understanding both plant trait evolution and plant responses to environmental change. While an observed negative SM-SO relationship is hypothesized to reflect universal constraints on resource allocation in all plants, domestication has likely fundamentally altered this relationship. Using a dataset of SM and SO for 41 of the world most widespread crops and 1,190 wild plant species, coupled with observational data on these traits in soy (*Glycine max*) and maize (*Zea mays*), I show that domestication has systematically rewired SM-SO relationships in crops. Compared to wild plants, virtually all crops express a higher SM for a given SO; this domestication signature is especially prominent in seed crops, and also influences the phylogenetic signal in SM and SO. In maize these traits have become positively related likely due to simultaneous selection for greater SM and SO, while in soy these traits have become decoupled likely due to primary selection for SM only. Evolved relationships between SM and SO in plants have been disrupted by both conscious and unconscious artificial selection, which represents a key aspect of how the functional biology of crops differ fundamentally from wild plants along “universal” plant trait spectra.

## Introduction

### Seed mass and seed output as a critical dimension of plant functional ecology

Differences in seed traits including mass, output, shape, dispersal, biochemical constitution, and dormancy have come to represent critical axes of life-history variation among plant species worldwide (Westoby et al. 1996, Moles et al. 2005, Diaz et al. 2016). Understanding and quantifying variation in these seed traits among terrestrial plant species has been central in better understanding a range of large-scale processes including the evolution of angiosperms and plant biogeography (e.g. Westoby et al. 1992, Thompson et al. 1993, Westoby et al. 1996, Moles and Westoby 2004, Moles et al. 2005), as well as multiple aspects of plant life history strategy including dispersal ability, stature, seedling competitiveness and survivorship, plant-and species responses to disturbance, colonization ability, and persistence of seeds in soils (Thompson et al. 1993, Westoby et al. 1996, Leishman et al. 2000, Moles and Westoby 2003, 2004). Of the seed traits explored in the comparative plant sciences, seed mass (SM) has received the most research attention (cf. Kattge et al. 2011), with comparative analyses including thousands of plant species indicating that SM varies by up to 6 orders of magnitude across species (Westoby et al. 2002, Moles and Westoby 2003). When considered independently or alongside other leaf-, stem- and whole-plant traits, variation in SM across species represents a key trait defining the functional ecology of plants worldwide (e.g. Westoby 1998, Cornelissen 1999, Westoby et al. 2002, Diaz et al. 2016).

In plant ecology, a long-recognized trade-off between SM and seed output (SO) remains central to plant resource allocation and life-history theories, particularly as it pertains to plant reproduction strategies (Harper et al. 1970, Smith and Fretwell 1974, Lloyd 1987, Westoby et al. 2002, Moles et al. 2004). Under conditions of finite resources, theory and observation suggest plant species differentiate from one another along a SM-SO axis that is hypothesized to optimize reproductive fitness and success, given a certain set of environmental conditions (Sadras 2007). In the simplest conceptual terms, the endpoints of the SM-SO trade-off are defined by plants allocating resources to a small number of large seeds, vs. plants that allocate resources to a large number of very small seeds; the wide variation that exists in between these conceptual points reflects “optimized” solutions to resource allocation (Sadras 2007). Similar to the evidence that supports the existence of universal trade-offs (or correlations) among other plant functional traits (e.g. Wright et al. 2004), global databases including trait values from thousands of species can be used to define a “universal SM-SO” trade-off that exists among plants globally. The shape of this trade-off reflects evolutionarily defined constraints on possible combinations of SM and SO that can (or are most likely to) occur across non-domesticated plant species (hereafter referred to as “wild plants”).

### Seed mass and seed output trade-offs and crops

Historical analyses(Meyer et al. 2012) coupled with recent observational and experimental studies, have refined our understanding of how plant trait variation and correlations have been fundamentally altered by crop domestication (Meyer et al. 2012, Milla et al. 2014, Martin et al. 2017). While the suites of traits that are under intentional and unintentional artificial selection is wide – including whole-plant, leaf-, and root traits (e.g. Milla et al. 2014) – plant yield components including SM and/ or SO have been under the most intensive selection (Sadras 2007).

Assuming that crops have been selected for increased SM and SO leads to the hypothesis (“Hypothesis 1) that, as compared to a “global SM-SO trade-off”, artificial selection results in all crops expressing a higher SM for a given SO (or vice versa) as compared to wild plants. Additionally though, artificial selection directly targets SM and SO only for a certain group of crops such as cereals (e.g. wheat, rice, and maize), oil seed crops, or legumes including soy and other pulses. Therefore, one may also hypothesize (“Hypothesis 2”) that for a given crop species, the degree of divergence away from a global SM-SO trade-off differs according to the plant organ under selection, with seed crops showing the strongest divergences on average.

Yet even for crops under selection for seeds, within-species SM-SO trade-offs may differ considerably according to plant growth form and reproductive strategy. In certain crops including maize – one of the crops employed in my analysis here – both increased SM (i.e. mean kernel mass) (Hufford et al. 2012) and SO (i.e. mean kernels per plant)(Brown et al. 2011) have been simultaneously targeted by artificial selection. This leads to the hypothesis (Hypothesis 3) that, counter to a global SM-SO trade-off, crops such as maize express a positive SM-SO correlation. Alternatively, genome sequencing indicates that increased SM in soybean – also employed in my analysis here – has been targeted during artificial selection (Liu et al. 2007, Zhou et al. 2015), while the number of inflorescences and ultimately SO remains plastic, and largely determined by local resource availability and concomitant plant growth rates (Andrade et al. 2005). This leads to the hypothesis (Hypothesis 4) that in soy, covariation along an intraspecific SM-SO trade-off would be weak or potentially non-existent. Here, I used a large dataset of SM and SO from 1,190 wild plant species and 41 of the world’s most widespread crops, derived from both functional trait databases and field studies on maize and soy, in order to test these four complementary hypotheses.

## Methods

### Generating a global SM-SO trade-off with functional trait data

Data for both SM and SO were acquired from a structured enquiry submitted to the TRY Functional Trait Database (Kattge et al. 2011). We specifically requested information on trait ID 26 (“seed dry mass”, corresponding to SM) and trait ID 131 (“seed number per plant”, corresponding to SO). This request returned *n*=117,882 and *n*=9,292 observations for SM and SO respectively. All statistical analyses were then performed using R version 3.4.0 (R Foundation for Statistical Computing, Vienna, Austria). First, for each observation I used the ‘TPL’ function in the “’Taxonstand’ R package (Cayuela et al. 2017) in order to cross-reference all species, genus, and family identities with The Plant List and resolve all synonyms or errors. Once taxonomy checks were complete, species-level mean values for both SM and SO were then calculated.

This list of species was then cross-referenced with a list of the world’s crop species reported by the Food and Agricultural Organization of the United Nations (FAO) (FAO 2018), which was refined to species-level taxonomy by Martin and Isaac (2015). This procedure led to identification of *n*=38 crop species which had both paired SM-SO data in TRY, and have been identified by the FAO as a commodity species. I therefore supplemented the TRY dataset with species mean SM and SO values extracted from the literature for a number of additional common crops including soy (*Glycine max(Hayes et al. 2018)*), sunflower (*Helianthus annuus*(Libenson et al. 2002), rice (*Oryza sativa*(Wang et al. 2008)), and maize (*Zea mays*(Maddonni and Otegui 2006)); data for SM and SO was also sought for all additional crops listed by Martin and Isaac (2015) which were not in TRY, but this data was not available in peer-reviewed literature. Therefore in sum, this data consolidation process resulted in paired SM-SO data for *n*=1231 species in total.

Each of the 41 crop species were then classified broadly according to the main commercial portion of the plant as one of: i) “seed plants” which are harvested for seeds (*n*=17 crop species); ii) “tree crops” which are harvested for timber or as ornamental species (*n*=3 crop species); iii) “other crops” which are harvested for other plant parts including leaves, roots, or large inflorescences (*n*=21 crop species); or iv) “wild species” which are non-domesticated plants (*n*=1190 species).

### Crops along an SM and SO trade-off

Analyses using the ‘fitdist’ function in the ‘fitdistrplus’ R package (Delignette-Muller and Dutang 2015) indicating that both the SM and SO datasets were better described by normal or log-normal distributions (as per lower log-likelihood scores; Table 1), so log-transformed data was used in all subsequent analyses. I first fit a standardized major axis regression (SMA) model to the entire log-SM and log-SO, in order to test for the presence of a “global SM-SO trade-off”. This SMA was implemented in using the ‘sma’ function in the ‘smatr’ R package(Warton et al. 2012), with 95% confidence limits surrounding the overall model generated through bootstrapping.

All SMA model residuals associated with each species were then extracted, and significant differences in residuals among the four different plant types were evaluated. Due to unequal sample sizes and non-independence of data points owing to phylogenetic structure in the data (see below), this test was performed as a linear mixed effects model using the ‘lme’ function in the ‘nlme’ R package(Pinheiro et al. 2016). Specifically, in this model SMA residuals were predicted as a function of plant type (as a fixed effect), while accounting for genus identity nested within family identity (as random effects). In addition to assessing overall significance of the plant type term (i.e. the fixed effect), I calculated mean (± S.E.) SMA residuals for each plant type using the ‘lsmeans’ function in the ‘lsmeans’ R package (Lenth 2016), and assessed the pairwise differences in mean SMA model residuals among all four plant types using a Tukey post-hoc tests (also implemented with the ‘lsmeans’ function). Post-hoc tests based on this linear mixed-effect model were considered significant when assessed against a Bonferonni-corrected *p*-value of 0.008.

### Phylogenetic signal in SM and SO

A phylogenetic tree was constructed for the entire *n*=1,231 species using Phylomatic (Webb and Donoghue 2005) to generate a phylogenetic tree, based on the Angiosperm Phylogeny Group megatree (“R20120829.new”). The BLADJ algorithm in Phylocom(Webb et al. 2008) was used to estimate phylogenetic branch lengths according to clade ages based on fossil records (Wikstrom et al. 2001) which were updated by Gastauer and Meira-Neto (2016). Unresolved evolutionary relationships were treated as polytomies.

Phylogenetic signal in log-SM and log-SO across the entire phylogeny was then quantified as Pagel’s *λ (Pagel 1999)*. For this analysis, a Pagel’s *λ* value equal to 0 represents instances of no phylogenetic signal (i.e. where evolution of SM and/or SO is entirely independent of phylogeny), and Pagel’s *λ* values of 1 represent instances where a phylogeny perfectly predicts trait data (i.e. where evolution of SM and/or SO perfectly matches a Brownian model of trait evolution) (Pagel 1999). Values of Pagel’s *λ* were calculated using the ‘phylosig’ function in the ‘phytools’ R package (Revell 2012). Significance tests for were performed as randomization tests (with *n*=1000 randomizations used), where SM and SO data were randomly shuffled across the phylogeny, and Pagel’s *λ* was recalculated on each randomized dataset; phylogenetic signal was considered statistically significant if the observed Pagel’s *λ* fell within the upper 95% of this randomized distribution.

In order to assess if the presence of crops influenced phylogenetic signal in SM and SO, I then recalculated Pagel’s *λ* with crop species removed from the dataset (where *n*=1,190 in this reduced phylogeny and trait dataset). Specifically, if Pagel’s *λ* was to increase when crops were removed from the phylogeny, this could be interpreted as crops reducing the strength of the phylogeny signal in SM or SO. All trait-phylogeny relationships were also graphed visually using the ‘plotTree.wBars’ function in the ‘phytools’ R package (Revell 2012).

### Soy and maize along an SM-SO trade-off

I employed two crop species-specific datasets, both derived from field studies, to evaluate if soy and maize expressed an SM-SO trade-off that was consistent with a global pattern (i.e. based on the *n*=1,231 species dataset). Soy data was taken from a published field study focused on soy leaf economics traits that was conducted on a 30-year old experimental farm in Guelph, Canada (43° 32′ N, 80° 12′ W) (Hayes et al. 2018). This study provided paired plant-level SM and SO data for *n*=45 soy plants (detailed in (Hayes et al. 2018)). Maize data was taken from two different field studies conducted at the same site, where paired SM and SO data was directly available (Maddonni and Otegui 2006, Mayer et al. 2012). Specifically, *n*=26 paired maize SM-SO observations were available in Table 1 of (Maddonni and Otegui 2006), while *n*=8 paired maize SM-SO data were taken from Table 1 of (Mayer et al. 2012). Data from both maize studies were derived from field experiments at the same site (the Experimental Station of the National Institute of Agricultural Technology) in Pergamino, Argentina (33° 56′ S, 60° 34′ W). I then used fit a SMA model (as described above) to both soy and maize datasets separately to describe intraspecific SM and SO patterns, and where SMA models were significant, 95% confidence limits were generated through bootstrapping with replacement (with 1000 replicates used) implemented with the ‘bootstrap’ function in the ‘modelr’ R package (Wickham 2017).

### Data availability

The compiled database of *n*=1,231 plant species used in my analyses here (presented in Figs. 1 and 4) are available as individual datasets in the TRY Functional Trait Database. Compiled data on soy and maize (presented in Fig. 3) are available upon request from the author, or in the original publications as cited in the methods.

**Fig. 1.**
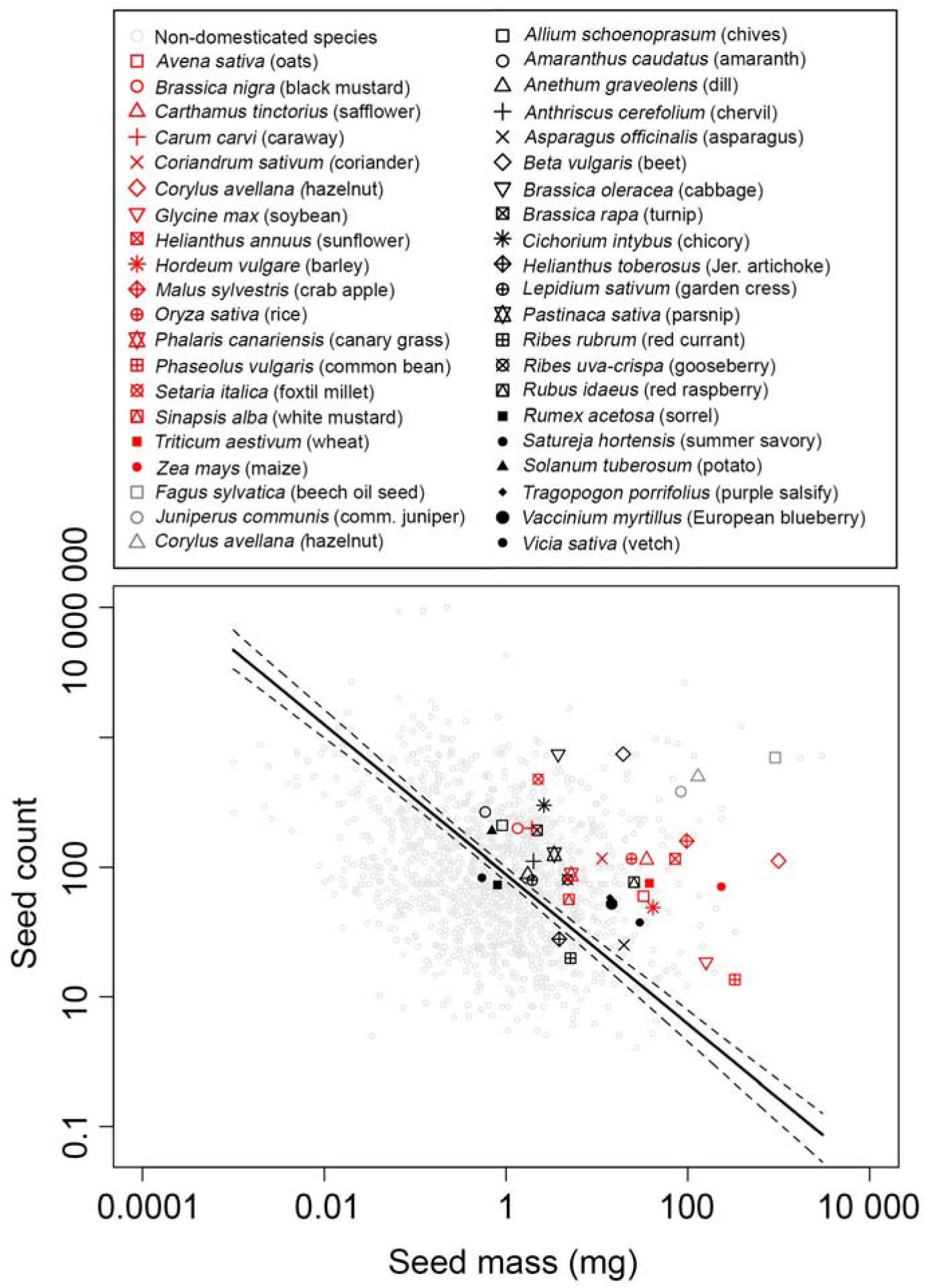
Relationship between seed mass and seed output across *n*=1190 non-domesticated species and *n*=41 of the world’s most widespread crops. Solid black trend line represents a standardized major axis regression fit across the entire dataset (*n*=1231) and dashed lines represent 95% confidence limits.

## Results

### Crops along a global seed mass seed output trade-off

Data from *n*=1231 species across 517 genera and 92 families demonstrate the presence of a SM-SO trade-off in plants globally, with a standardized major axis (SMA) regression slope of −0.86 (95% C.I. = −0.91, −0.82, SMA *r*^2^=0.06, *p*<0.0001; Fig. 1). Of the 41 crop species represented in the SM-SO dataset, 90.2% (37 crop species) feel above the global SM-SO axis (where SMA model residuals ≥ 0) while only four fell below the primary SM-SO axis (Fig. 1). In comparison, 46.1% and 53.9% of the *n*=1190 wild plant species were approximately evenly distributed above and below the primary SM-SO model, respectively (Figs. 1 and 2).

**Fig. 2.**
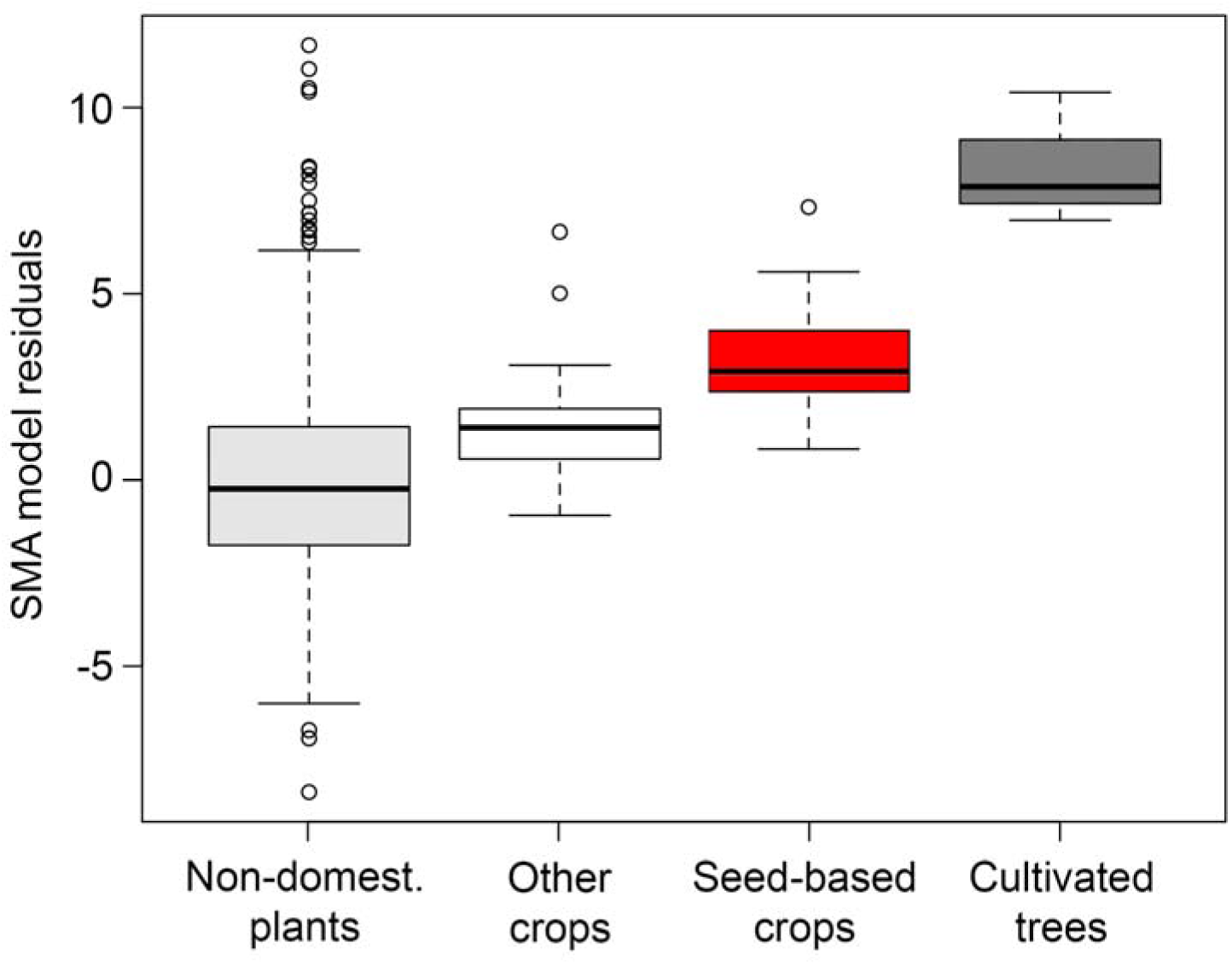
Deviations from a global seed mass-seed output trade-off in three types of crops and non-domesticated plants. Deviations from a global SM-SO relationship are calculated as residuals from a standardized major axis regression fit to *n*=1231 plant species (see Fig. 1). “Other crops” (*n*=21) correspond to plant species cultivated for vegetative-or belowground structures. Sample sizes for seed-based crops, cultivated trees, and non-domesticated plants are *n*=17, *n*=3, and *n*=1190, respectively.

Across all 1231 species, SMA model residuals ranged from −8.4 to 11.7 (average residual=1.14e-11±2.5 (S.D.)). While accounting for the phylogenetic non-independence of data (Figs. S1 and S2) and uneven sample sizes, the extent of divergence from the SM-SO axis differed significantly across crop types (mixed-effects model *F*_4, 711_=13.1, *p*<0.001, Fig. 2). Specifically, SMA residuals for seed- and tree crops were significantly greater than zero (*p*≤ 0.001), while residuals for non-seed/ tree crops and wild plant species did not differ from zero (*p*≥0.067, Table S1).

There was a gradation of divergence from a central SM-SO trade-off (Fig. 2). Tree crops and seed crops expressed the highest SM for a given SO, and in comparison non-seed crops and wild plant expressed lower SM for a given SO; these groups broadly differed significantly from one another in terms of average divergence from a central SM-SO axis (Table S1). Crops harvested for seeds diverged significantly more strongly from a central SM-SO trade-off as compared to wild plants (post-hoc contrast *p*≤0.0001; Table S2). Furthermore, 17 seed crops expressed higher average SMA residuals (3.4±1.6 (S.D.), in comparison to the average SMA residuals observed in 21 non-seed crops (1.6±1.8 (S.D.) (Fig. 2, Table S2) The three tree crops in the dataset differed most strongly from a central SM-SO trade-off, showing SMA residuals which were significantly higher than all other crops and wild plants (8.4±1.8 (S.D.); post-hoc *p*≤0.04 for all three contrasts; Fig. 2). Among seed crops, notable divergences from the global SM-SO model included hazelnut (*Corylus avellana*, SMA residual=7.3), maize (*Zea maize*, SMA residual=5.1), and sunflower (*Helianthus annuus*, SMA residual=4.8).

### Within-crop seed mass seed output trade-offs

Data from soy (*n*=45) and maize (*n*=35) indicated that SM-SO patterns differed both between these crops, as well as in comparison to a global SM-SO pattern. In maize SM and SO covaried significantly among individual plants (SMA *r*^2^=0.334, *p*<0.0001; Fig. 3), however artificial selection has resulted in this relationship being positive and therefore opposite that of a global SM-SO pattern (maize SMA slope=0.27, 95% C.I. = 0.20, 0.37, Fig. 3). Alternatively, soy data indicate that targeted selection for higher SM alone has resulted in a decoupling of SM and SO (Fig. 3). These two seed traits were not significantly related in soy, expressing only a weak negative relationship (SMA slope=-0.14, 95% C.I.=-0.2, −0.11, SMA *r*^2^=0.002, *p*=0.757; Fig. 3). Consistent with reproductive allocation theory and targeted selection for SM(Sadras 2007), the lack of a significant SM-SO relationship in soy was qualitatively associated with variation in SM which was nearly an order of magnitude lower than variation in SO (where CV=9.5 and 76.3, respectively; Table S1).

**Fig. 3.**
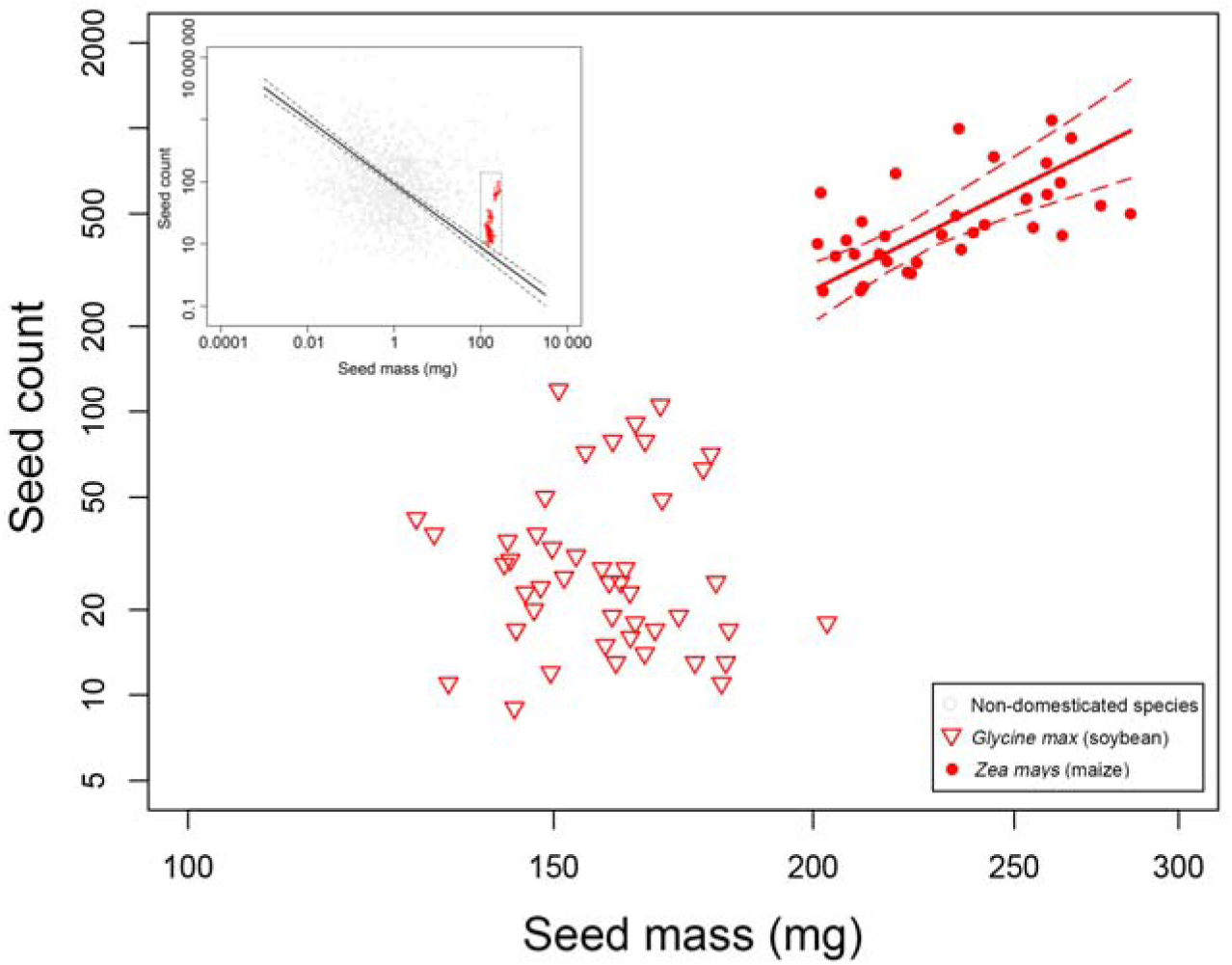
Seed mass and seed output in soybean (*Glycine max*, *n*=45) and maize (*Zea maize*, *n*=37). Data are presented in relation to a SM-SO trade-off observed across 1231 plant species (gray rectangle, inset graph). Solid red trend line represents a standardized major axis (SMA) regression fit to the maize data with red dashed lines representing 95% confidence limits. The SMA model fit to soy data was not significant, so is not presented here.

### Phylogenetic signal along the seed mass-seed output trade-off

Across the entire dataset (*n*=1231) both SM and SO expressed significant phylogenetic signal, with Pagel’s *λ* of 0.995 and 0.853, respectively (*p*<0.001 in both cases, Fig. 4). When crops are removed from the dataset, SM and SO are better predicted by phylogenetic relatedness than when artificially crops are present. Specifically, excluding crops from the datasets resulted in a small, albeit detectable, increase in Pagel’s *λ* for both SM (0.996 with crops) and SO (0.859 with crops).

**Fig. 4.**
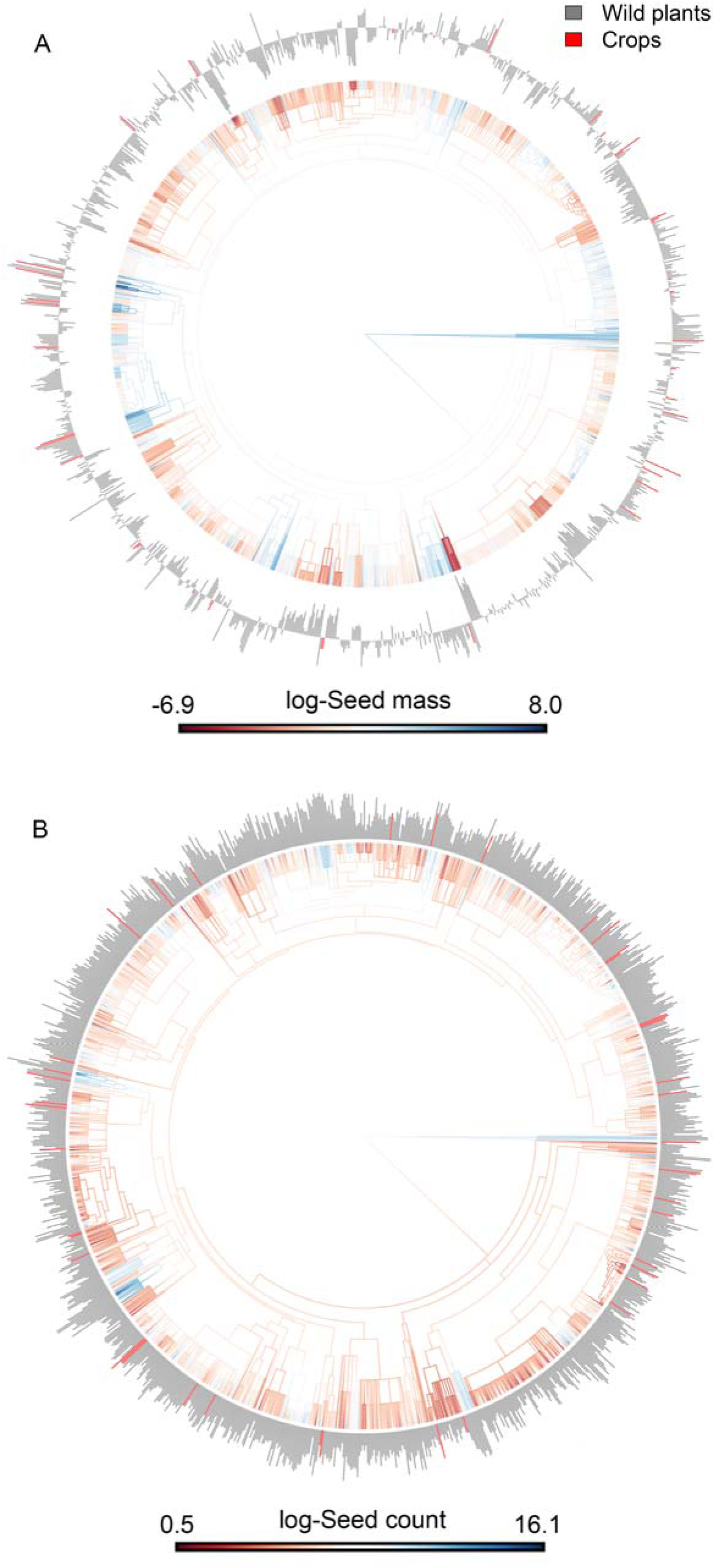
Variation in seed mass (A) and seed output (B) across a phylogeny of 1,231 crop- and wild plant species. Bars across the tips of the phylogeny represent mean seed mass and seed count (plant^-1^) values, with crop species highlighted in red and wild plants in gray.

## Discussion

### The functional profile of crops vs. wild plants

Research documenting the traits that have been targeted by crop domestication suggests that SM and/or SO are among those under the most intensive artificial selection (Meyer et al. 2012). While much of this research has taken archaeological and/or genetic approaches the analysis presented here refines a functional trait-based signature of crop domestication and artificial selection. Specifically, consistent with my hypotheses i) artificial selection has led to most crops significantly deviating from a global SM-SO trade-off (Fig. 1), such that ii) crops under selection for seeds deviate most strongly as compared to non-seed crops (Fig. 2). Yet within seed crops the degree to which artificial selection has reshaped the evolved biomechanical and/ or physiological constraints on trait syndromes (or trait trade-offs), depends on the specific reproductive traits/ yield components being targeted (Fig. 3). In certain crops such as maize, artificial selection has fundamentally altered the direction of trait trade-offs imposed by natural selection, while in others such as soy artificial selection may only act to decouple the covariation among reproductive traits (Fig. 3)

When taken with studies examining other suites of crop functional traits and trait spectra (Milla et al. 2014, Martin et al. 2017), the shifts in SM-SO trade-offs in crops vs. wild plants observed here (Figs. 1-3) represent components of a broader “disruption” in plant resource-use syndromes incurred by artificial selection (Milla et al. 2014). Other and observational data indicate that crops generally express greater values of resource capture traits as compared to their wild progenitors. Indeed, detailed analyses of leaf traits indicate that certain crops express among the most extreme “resource-acquiring” trait syndromes in plants globally (Martin et al. 2018), while at the same time artificial selection has resulted in trait relationships, or “phenotypic integration”(Milla et al. 2014), that are considerably weaker in crops vs. wild plants (Milla et al. 2014, Martin et al. 2017).

While neither SM nor SO values in crops have been shifted to extreme ends of a reproduction trait spectrum (Fig. 4), the analysis here indicates that relationships between SM and SO within individual species may be fundamentally rewired through domestication (Fig. 3). Our results and other studies exploring trait syndrome disruption (e.g. Milla et al. 2014) provide compelling evidence to indicate crops fundamentally differ from wild plants along a global spectrum of plant form and function(Diaz et al. 2016). Quantifying the position of crops and how their traits trade-off along a global trait spectra (Wright et al. 2004, Diaz et al. 2016) represents a means of synthetically defining the functional ecology of crops, which in turn would support a range of hypotheses on the unintended impacts of artificial selection.

### Conscious and unconscious selection and trait trade-offs in crops

Studies employing quantitative trait locus (QTL) mapping would suggest that the results here deviations of crops away from a central SM-SO trade-off (Fig. 1), have likely occurred largely in response to “conscious selection” for these individual traits (e.g. Tao et al. 2017). Major shifts in the shape of the SM-SO relationship in the two crops explored here is also consistent with strong conscious selection for these particular traits (Fig. 3). Furthermore, analyses of phylogenetic patterns in seed traits here demonstrated a small albeit detectable reduction in phylogenetic signal when crops are removed from analyses (Fig. 4). Major deviations in SM and SO among crops vs. wild relatives, both here and in experimental studies, are most likely consistent with long-term conscious selection of seed traits.

At the same time though, theories from functional trait-based ecology do hypothesize that unconscious selection has also played a role in reshaping SM-SO relationships in crops. Specifically, from a plant resource allocation/ natural selection perspective, higher SM at a given SO in open cultivated agricultural environments would be expected to competitive benefits to crop plants, including greater seedling growth rates and survival at greater burial depths. Yet while this theory derived from plant resource allocation theory suggests that unconscious selection may also have contributed to the patterns observed here, experimental evidence in support of this hypothesis is currently lacking (reviewed by Milla et al. 2015). Detailed partitioning of the relative importance of conscious vs. unconscious selection remains a leading avenue for better understanding the genetic vs. phenotypic controls on SM, SO, and their relationships in crops.

